# A Convolutional Network Architecture Driven by Mouse Neuroanatomical Data

**DOI:** 10.1101/2020.10.23.353151

**Authors:** Jianghong Shi, Michael A. Buice, Eric Shea-Brown, Stefan Mihalas, Bryan Tripp

## Abstract

Convolutional neural networks trained on object recognition derive some inspiration from the neuroscience of the visual system in primates, and have been used as models of the feedforward computation performed in the primate ventral stream. In contrast to the hierarchical organization of primates, the visual system of the mouse has flatter hierarchy. Since mice are capable of visually guided behavior, this raises questions about the role of architecture in neural computation. In this work, we introduce a framework for building a biologically constrained convolutional neural network model of lateral areas of the mouse visual cortex. The structural parameters of the network are derived from experimental measurements, specifically estimates of numbers of neurons in each area and cortical layer, the interareal connectome, and the statistics of connections between cortical layers. This network is constructed to support detailed task-optimized models of mouse visual cortex, with neural populations that can be compared to specific corresponding populations in the mouse brain. The code is freely available to support such research.

## Introduction

Convolutional neural networks (CNNs) trained on object recognition derive some inspiration from the neuroscience of the visual system in primates, and have been used as models of the feedforward computation performed in the primate ventral stream (1–3). Indeed, these networks show functional similarity to areas of the primate visual system (4, 5) and the task-training paradigm for artificial neural networks is known to correlate with more predictive models and more similar representations to neural activity in both visual and auditory areas (6–9). Despite these successes and the clear power of CNNs to solve machine learning problems in the visual domain (10, 11), they are not precise analogues for the biological circuit. Recent endeavors (12, 13) show that imposing brain like structure such as shallowness and recurrence can improve the functional similarity to the primates’ brain. The interplay of architecture and computation remains an open problem in both machine learning and neuroscience.

In contrast to the hierarchical organization of primates, the visual system of the mouse has a shallower organization in which multiple higher visual areas are at similar levels of the hierarchy (14). Since mice and other rodents are capable of visually guided behavior including invariant object recognition (15, 16), this raises questions about the role of architecture in neural computation.

In the mean time, large scale tract tracing data sets have revealed neuro-anatomical structure in unprecedented detail (17–19). Using such data sets, we construct detailed estimates for circuit properties of the mouse visual system, including the interarreal connectome and the numbers of neurons per region and layer.

Convolutional networks share weights across the visual field, and thus form an approximation of the likely functional properties of the real neural system, an approximation likely partially imposed by a degree of translation invariance of stimuli in the natural world leading to equivariant representations in neural systems (1–3). This weight sharing makes them much easier to train, which is an important practical consideration for establishing representations without a lifetime of experience. Python code for this project is available at https://github.com/mabuice/Mouse_CNN.

## Methods

### Construction of the MouseNet CNN

In this section, we introduce our framework for constructing the MouseNet CNN. Fig 1 shows an overview of this framework. The basic idea is to use available sources of anatomical data (e.g. tract tracing data, cell counts, and statistics of shortrange connections) to constrain the CNN network structure and architectural hyperparameters. We discuss the details of each step below.

**Fig. 1.**
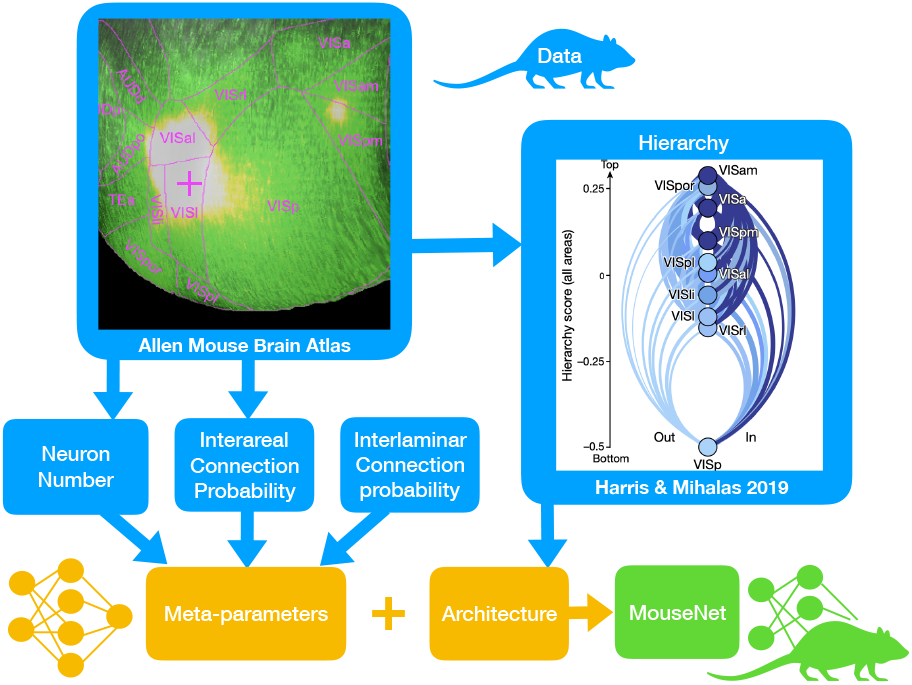
Modeling framework. Framework for constructing biologically constrained MouseNet.

#### Network architecture

MouseNet spans the ventral lateral geniculate nucleus (LGNv) and six visual areas (Fig 2A). Input to the network passes first through LGNv, and then to the primary visual area VISp. After VISp, the architecture branches into five parallel pathways, representing five lateral visual areas: VISl (lateral visual area), VISal (anterolateral), VISpl (posterolateral), VISli (laterointermediate), and VISrl (rostrolateral). Finally, the output of VISp together with all five lateral visual areas provide input to VISpor (postrhinal). We include only the lateral areas because they are more associated to object recognition while the medial areas are more involved in multimodal integration (20). The three-level architecture among the VIS areas was derived from an analysis of the hierarchy of mouse cortical and thalamic areas (Figure 6e in (14)), which considered feedforward and feedback connections from each area. In this analysis, VISp was clearly low in the hierarchy, and VISpor was clearly high, but the other lateral visual areas had similar intermediate positions.

**Fig. 2.**
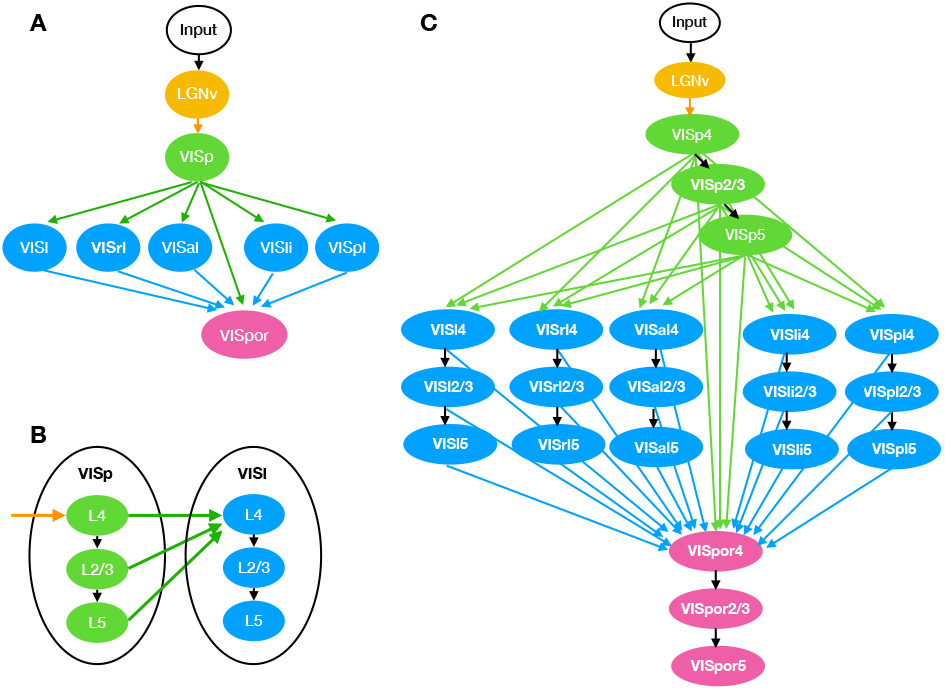
Illustration of MouseNet architecture. Only feedforward connections are included. (A) High-level organization of MouseNet, based on analysis of the hierarchy of lateral visual areas ((14)). (B) Connection patterns at the level of cortical layers. (C) Full MouseNet architecture.

In the MouseNet model, each VIS area is represented by three separate cortical layers: layer 4 (L4), layer 2/3 (L2/3) and layer 5 (L5). We call a specific cortical layer within a specific area a “region”. Here we only consider the feedforward pathway, thought in primates to drive responses within ≈ 100ms of stimulus presentation (2, 6). Following depictions of the canonical microcircuit (e.g. as summarized in Fig 5 in (21)), we consider the feedforward pathway to consist of laminar connections from L4 to L2/3, and from L2/3 to L5. Input from other areas feed into L4 and all of L4, L2/3 and L5 output to downstream areas, as shown in Fig 2B. This is consistent with broad connectivity among visual areas from each of these layers (Figure 2f of (14)). Fig 2C shows the MouseNet architecture in full detail, including all 22 regions and associated connections.

#### From architecture to convolutional neural net

Similar to the CALC model for the primate visual cortex by one of the authors (22), the general idea is to use convolution (Conv) operations to model the projections between different regions in the mouse visual cortex. Conv operations are linear combinations of many inputs, so they impose the assumption of linear synaptic integration. They are widely used in machine learning, because they run efficiently on graphical processing units, and they share parameters across the visual field, leading to reduced memory requirements and faster learning, relative to general linear maps. Each connection from source brain region *i* to target brain region *j* is modelled with a Conv operation, Conv^*ij*^. The input to Conv^*ij*^ corresponds to the neural activities in source region *i*, and the output of Conv^*ij*^ drives neural activities in the target region *j*. For example, as shown in Fig 3A, the projection from Region 1 to Region 2 (Proj 1→2) is modeled by Conv^12^. The neural activities in Region 1 correspond to the input to Conv^12^, while the neural activities in Region 2 are a nonlinear function (ReLU, as shown in Fig 3C) of the output of Conv^12^. In MouseNet, L4 of all areas except VISp receive multiple converging inputs, similar to Region 4 in the figure. In these cases, each projection is modeled by a separate Conv layer, and a nonlinear function (ReLU) is applied to the sum of the output from all these Conv layers, to produce the neural activities in Region 4.

**Fig. 3.**
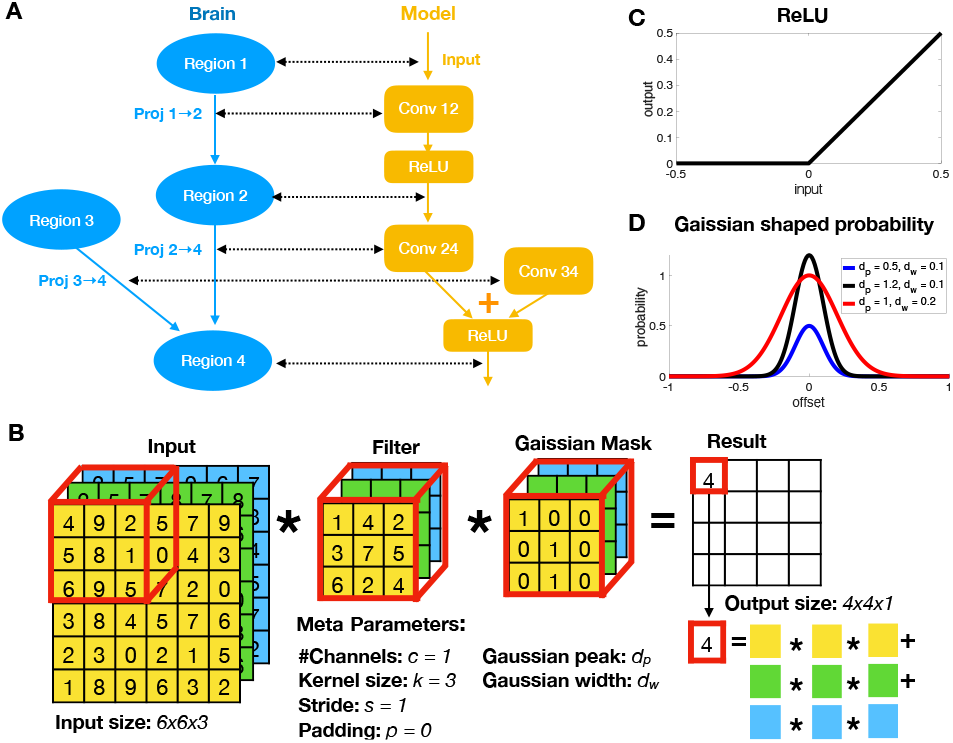
From mouse brain to CNN model. (A) From mouse brain hierarchy to CNN architecture. (B) An example of Conv operation with Gaussian mask. (C) ReLU operation in the CNN architecture. (D)Gaussian shaped probability for Gaussian mask.

#### Finding meta-parameters consistent with mouse data

After fixing the architecture, we need to determine the metaparameters for constructing the kernels for each Conv operation (Fig 3). The standard Conv operation is defined in terms of a four-dimensional kernel. The output of the kernel is a three-dimensional tensor of activations, *A^j^*, which pass through element-wise ReLU nonlinearities to produce non-negative spike rates. Element 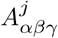 is the activation of the neuron at the *α^th^* vertical and *β^th^* horizontal position in the visual field, in the *γ^th^* channel (or feature map). The *γ^th^* channel of the activation tensor for region *j*, 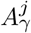, depends on inbound connections as,

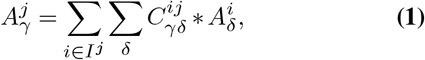

where *I^j^* is the set of regions that provide input to region *j*. Note that both 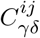 and 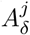 are two-dimensional, and they undergo standard two-dimensional convolution. The metaparameters of kernel *C^ij^* are: number of input channels 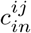, number of output channels 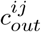, stride *s^ij^*, padding *p^ij^*, and finally kernel size *k^ij^*, i.e. the height and width (which are set equal) of 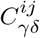. To make the connections realistically sparse, we add a binary Gaussian mask on the Conv filters, whose parameters are also estimated from data. See Fig 3B for an illustration of Conv operation with Gaussian mask. We constrain these meta-parameters with quantitative data whenever possible, and reasonable assumptions indicated by experimental observations otherwise.

#### Cortical population constraints

We set the horizontal and vertical resolution of the input (in pixels) based on mouse visual acuity. According to (23), the upper bound for visual acuity in mice is 0.5 cycles/degree, corresponding to a Nyquist sampling rate of 2 pixels/cycle x 0.5 cycles/degree = 1.0 pixel/degree. The visual stimulus covers about 112 degrees of visual angle in the horizontal direction and 64 degrees in the vertical direction, but we simplified this to square input of size 64 by 64 pixels.

The resolution of the other regions depends on both the resolution of the input, and strides of the connections. The stride of a connection is the sampling period with respect to its input. For example, a Conv with a stride of one samples every element of its input, whereas a Conv with a stride of two samples every other element (both horizontally and vertically), leading to output of half the size in each dimension. Because cortical neurons are not organized into discrete channels in the same way as convolutional network layers, there is no strong anatomical constraint on the stride. However, the mean stride has to be somewhat less than two; there are ten steps in the longest path through MouseNet, but if only six of them had a stride of two, the 64×64 input would be reduced to 1×1 in VISpor, with no remaining topography. Lacking strong constraints, for simplicity, we first attempted to set all the strides to one, but this left very few channels in some of the smaller regions (due to an interaction between channels and strides that we describe below). We therefore set the strides of the connections outbound from VISp to two, and others to one. The feature maps of LGNv and VISp were therefore 64×64 (the same as the input), and all others were 32×32.

Given the resolutions of the channels in each region, the numbers of channels are constrained by the number of neurons. Specifically, Let *n^i^* be the number of neurons in region *i* and 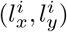 be the size of the output in area *i*, then the number of channels in area *i* is determined by 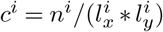.

#### Estimating number of neurons in each area from data

We only model the excitatory neural population in our model, consistent with the fact that all neurons in the model project to other visual areas. In fact, neurons in convolutional networks are neither excitatory or inhibitory, but have both positive and negative output weights. However, past modelling work (24, 25) has shown that such mixed-weight projections can be transformed so that the original neurons are all excitatory, and an additional population of inhibitory neurons recovers the functional effects of the original weights. We used estimates of the numbers of excitatory neurons in LGNv, VISp, VISal, VISam, VISl, VISpl, and VISpm from (26) (summarized in Tabel 1).

**Table 1.**
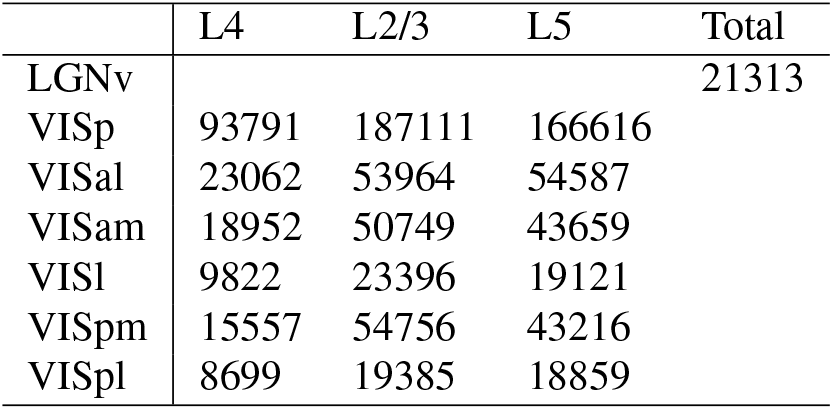
Exitatory population number given by (26).

|  | L4 | L2/3 | L5 | Total |
| --- | --- | --- | --- | --- |
| LGNv |  |  |  | 21313 |
| VISp | 93791 | 187111 | 166616 |  |
| VISal | 23062 | 53964 | 54587 |  |
| VISam | 18952 | 50749 | 43659 |  |
| VISl | 9822 | 23396 | 19121 |  |
| VISpm | 15557 | 54756 | 43216 |  |
| VISpl | 8699 | 19385 | 18859 |  |

For VISrl, VISli, and VISpor, we use the estimation of surface area from the voxel model (see section, *Estimating 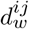 and 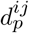 for interareal connections,* for details) and estimate their numbers with respect to VISl, assuming these areas have similar neural density. The estimated surface areas with population numbers are given in Table 2.

**Table 2.** Exitatory population number estimated by voxel model with respect to VISl in (26).

|  | surface area | L4 | L2/3 | L5 |
| --- | --- | --- | --- | --- |
| VISl | 0.92791 | 9822 | 23396 | 19121 |
| VISrl | 0.69805 | 7389 | 17600 | 14384 |
| VISli | 0.43561 | 4611 | 10983 | 8976 |
| VISpor | 1.39366 | 14752 | 35139 | 28719 |

#### Cortical connection constraints

Neurons tend to receive relatively dense inputs from other neurons that are above or below them, in other cortical layers, and the connection density decreases with increasing horizontal distance. Similarly, inputs from other cortical areas tend to have a point of highest density, with smoothly decreasing density around that point. We approximate such connection-density profiles with twodimensional Gaussian functions. Specifically, we model the fan-in connection probability from source region *i* to target region *j* at position (*x, y*) (position offset from center in *μm*) as,

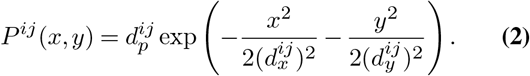

For simplicity, we assume 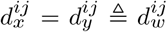 and 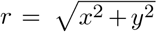 denote the offset from the center of the source layer, the above equation then simplifies to,

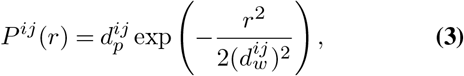

where 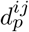 is the peak probability at the center and 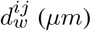 is the Gaussian width.

Both 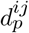 and 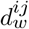 are estimated from mouse data. The parameters for interlaminar connections are estimated from hit-rate probabilities in functional connection studies, and the parameters for interareal connections are estimated from the mouse connectome (details below).

#### Conv layer with Gaussian mask

To translate our Gaussian models of connection density into network meta-parameters, we apply a binary mask to the weights of the Conv layers (Fig 3B). To do that, we first change the unit of 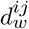 in Eq.4 from micrometers to source area-dependent “pixels” (unit of output size of source area *i*) by multiplying it with 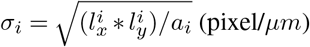. Here *a_i_* denotes the surface size of area *i*, estimated from the voxel model (See section, *Estimating 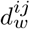 and 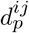 for interareal connections).* We then have,

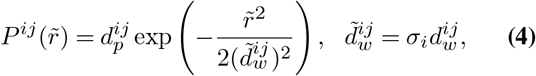

where both 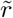 and 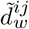 are in the “pixel” unit. The kernel size of the Conv layer is set to be 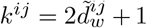, with padding calculated by 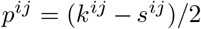. During initialization, a mask containing zeros and ones is generated for each Conv layer, with size 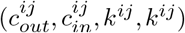. The probability of each element being one is 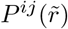, where 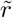 (pixel) is the offset from the center of mask. The weights of the Conv layer are then multiplied by the mask. This gives the connections realistic densities (or sparsities), with realistic spatial profiles.

#### *Estimating* 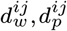 for interlaminar connections

For the interlaminar connections, we estimate the Gaussian width 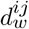 from multiple experimental resources. Firstly, from Table 3 in (27), we get the estimation of 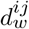 to be 114 micrometers for functional connections between pairs of L4 pyramidal cells in mouse auditory cortex. Secondly, manually extracted from (28) Fig 8B, we obtain the variation of the Gaussian width depending on source and target layer from cat V1. Finally, we use this variation to scale the L4 to L4 width 114 *μm* to other layers in the mouse cortex. We summarize the Gaussian widths from cat cortex, along with corresponding scaled estimates for mouse cortex, in Table 3. Note that in the current model, we only use the values for connections from L4 to L2/3 and from L2/3 to L5 (Fig 2B).

**Table 3.** Estimated Gaussian width 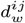 for interlaminar excitatory connections. The values outside of the parenthesis are extracted from (28); the values inside the parenthesis are scaled to mouse cortex, using the width 114 *μm* for L4-to-L4 connections in mouse auditory cortex (27). Units are micrometers (*μm*).

|  |  | Target |  |  |
| --- | --- | --- | --- | --- |
|  |  | L2/3 (scaled) | L4 (scaled) | L5 (scaled) |
| Source | L2/3 | 225 (142.5) | 50 (31.67) | 100 (63.33) |
|  | L4 | 220 (139.33) | 180 (114) | 140 (88.67) |
|  | L5 | 150 (95) | 100 (63.33) | 210 (133) |

To estimate the Gaussian peak probability 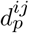, we first collect the connection probability between excitatory populations offset at 75 micrometer 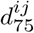 (Figure 4A in (29)). We then get the peak probability 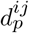 by the relation

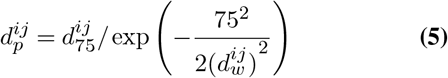

We summarize the probability at 75 micrometers 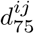 along with the peak probability 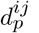 in Table 4.

**Table 4.**
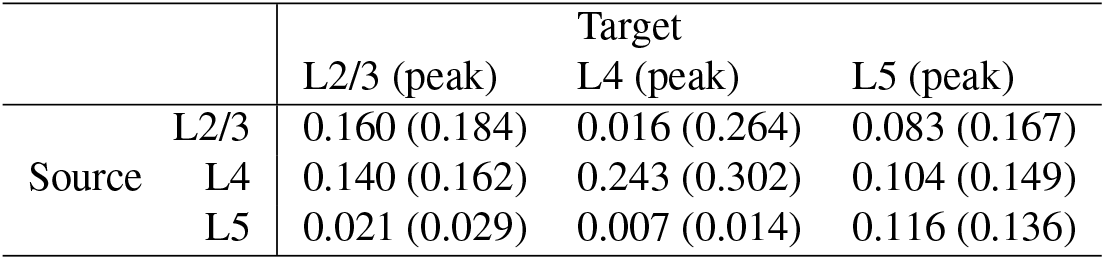
The connection probability between excitatory populations offset at 75 micrometer 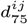. The numbers are from Fig 4A in (29). The calculated Gaussian peak probability 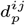 are given in parenthesis.

|  |  | Target |  |  |
| --- | --- | --- | --- | --- |
|  |  | L2/3 (peak) | L4 (peak) | L5 (peak) |
| Source | L2/3 | 0.160 (0.184) | 0.016 (0.264) | 0.083 (0.167) |
|  | L4 | 0.140 (0.162) | 0.243 (0.302) | 0.104 (0.149) |
|  | L5 | 0.021 (0.029) | 0.007 (0.014) | 0.116 (0.136) |

#### Estimating 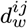 and 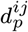 for interareal connections

To estimate interareal connection strengths and spatial profiles, we use the mesoscale model of the mouse connectome (19, 30). This model estimates connection strengths between 100 micrometer resolution voxels, based on 428 individual anterograde tracing experiments mapping fluorescent labeled neuronal projections in wild type C57BL/6J mice.

#### Flat map

The voxel model is in 3 dimensional space. For the purpose of our analysis, we need to map the 3 dimensional structure into 2 dimensions. First, we fit the visual area positions by a sphere with center 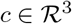 and radius *r*. Each position 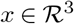 is then mapped to 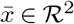 with relation

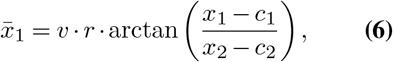

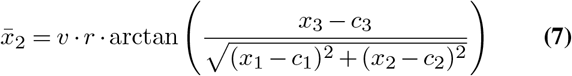

where *v* = 100*μm* is the size of the voxel.

#### Area size

Estimates of the size of each area/layer are needed to convert the widths of connection profiles from voxels in the mesoscale model to convolutional-layer pixels in MouseNet. For this purpose, each area’s size is approximated by the area of a convex hull of its mapped two-dimensional positions. These estimates are summarized in Table 5.

**Table 5.** Area size (*mm*^2^) estimated from the voxel model.

|  | VISp | VISal | VISl | VISli | VISpl | VISrl | VISpor |
| --- | --- | --- | --- | --- | --- | --- | --- |
| L4 | 4.3271 | 0.4909 | 0.8793 | 0.3355 | 0.2865 | 0.6182 | 0.5264 |
| L2/3 | 4.7406 | 0.5477 | 0.9279 | 0.4356 | 0.6659 | 0.6980 | 1.3937 |
| L5 | 4.2511 | 0.4972 | 0.8651 | 0.4039 | 0.6785 | 0.6748 | 1.2445 |

#### Estimating 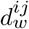

For each connection from source region *i* to target region *j*, we estimate 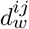 from the mesoscale model. The first step is to estimate the widths of connections to individual voxels in *j*. The incoming width 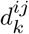 for target voxel *k* in *j* is estimated by the standard deviation of the connection strength about its center of mass. Specifically, 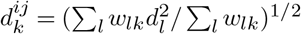, where *l* indexes the voxels in source region *j*, *w_lk_* is the connection weight between source and target voxels *l* and *k* in the mesoscale model, and *d_l_* is the distance from voxel *l* to the center of mass of these connection weights. We then estimate 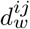 as the average of these widths over the voxels in *j*. We omit from this average any target voxels that have multi-modal input profiles. This procedure provides an upper bound for 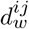, because a target voxel may include multiple neurons with partially overlapping input areas.

#### Estimating 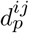

The mesoscale model provides estimates of relative densities of connections between pairs of voxels. But an additional factor is needed to convert these relative densities into neuron-to-neuron connection probabilities. For this purpose, we assumed that each neuron received inputs from 1000 neurons in other areas (we call this number the extrinsic in-degree, *e*). This is on the order of the estimate from Fig S9 M in (31). Given this assumption, we calculated 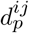 by the relation,

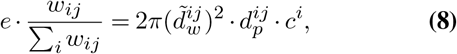

where *w_ij_* is the connection strength from source area *i* to target area *j*, estimated from integrating the connection weights of the corresponding areas in the mesoscale model. The estimated values for 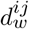 and 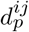 are given in Table 7.

**Table 6.** Parameters from data or assumptions

| Notation | CNN parameter | Biological source or assumptions |
| --- | --- | --- |
| $n^i$ | Number of neurons in area $i$ | Based on (26) combined with the voxel model (19) |
| $a_i$ | 2 dimensional area size for area $i$ | Estimated from voxel model data (19) |
| $e$ | Total fan-in connections for all areas | Set to be 1000 based on (31) |
| $(l_x^0, l_y^0)$ | input size | 64x64 based on mouse visual acuity (23) |
| $(l_x^i, l_y^i)$ | output size of area $i$ | 64x64 up to VISp, 32x32 after VISp (Assumption) |
| $d_w^{ij}$ | Gaussian width (interlaminar) | From mouse (27) and cat (28) data |
|  | Gaussian width (interareal) | Estimated from voxel model (19) |
| $d_p^{ij}$ | Gaussian peak (interlaminar) | Based on hit rate experiments (29) |
|  | Gaussian peak (interareal) | Estimated from voxel model (19) |

**Table 7.**
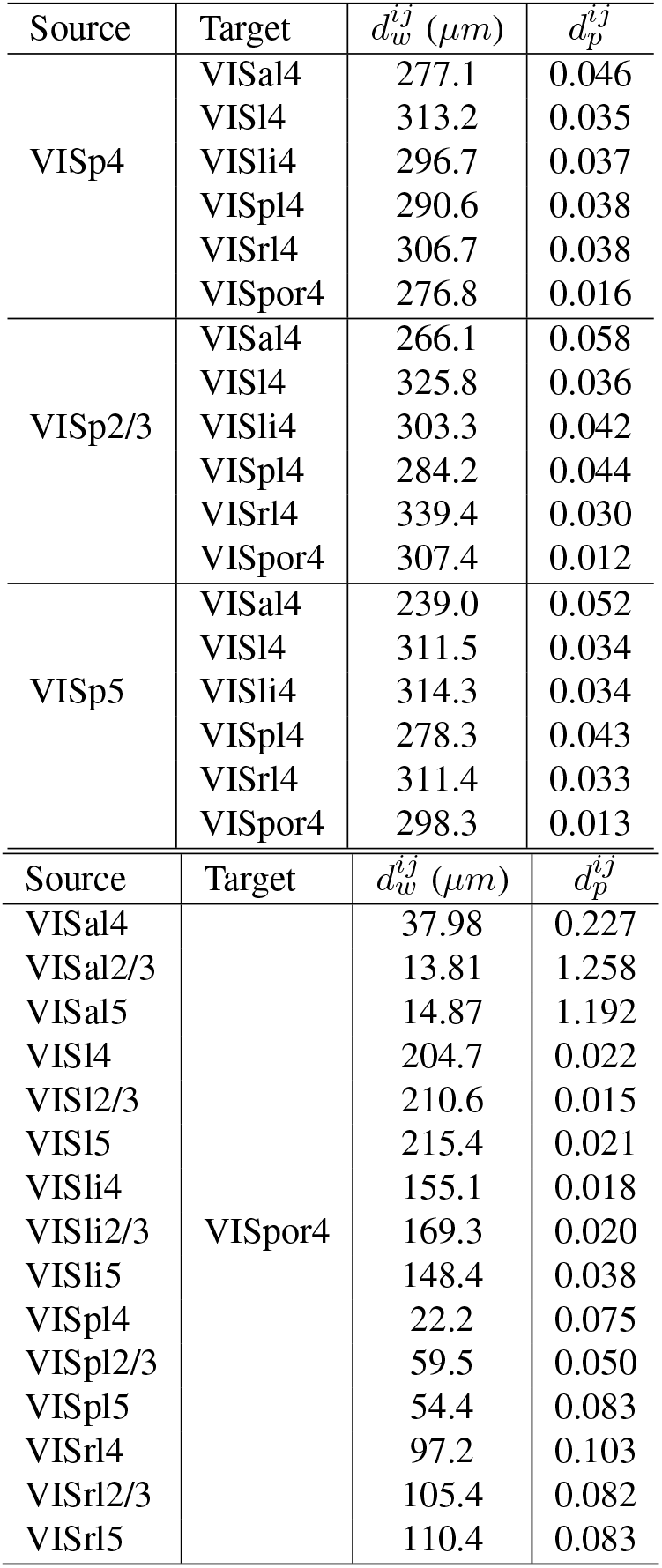
The estimated 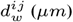 and 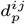 for interareal connections.

| Source | Target | $d_w^{ij}$ ($\mu m$) | $d_p^{ij}$ |
| --- | --- | --- | --- |
| VISp4 | VISal4 | 277.1 | 0.046 |
|  | VISl4 | 313.2 | 0.035 |
|  | VISli4 | 296.7 | 0.037 |
|  | VISpl4 | 290.6 | 0.038 |
|  | VISrl4 | 306.7 | 0.038 |
|  | VISpor4 | 276.8 | 0.016 |
| VISp2/3 | VISal4 | 266.1 | 0.058 |
|  | VISl4 | 325.8 | 0.036 |
|  | VISli4 | 303.3 | 0.042 |
|  | VISpl4 | 284.2 | 0.044 |
|  | VISrl4 | 339.4 | 0.030 |
|  | VISpor4 | 307.4 | 0.012 |
| VISp5 | VISal4 | 239.0 | 0.052 |
|  | VISl4 | 311.5 | 0.034 |
|  | VISli4 | 314.3 | 0.034 |
|  | VISpl4 | 278.3 | 0.043 |
|  | VISrl4 | 311.4 | 0.033 |
|  | VISpor4 | 298.3 | 0.013 |
| Source | Target | $d_w^{ij}$ ($\mu m$) | $d_p^{ij}$ |
| VISal4 | VISpor4 | 37.98 | 0.227 |
| VISal2/3 |  | 13.81 | 1.258 |
| VISal5 |  | 14.87 | 1.192 |
| VISl4 |  | 204.7 | 0.022 |
| VISl2/3 |  | 210.6 | 0.015 |
| VISl5 |  | 215.4 | 0.021 |
| VISli4 |  | 155.1 | 0.018 |
| VISli2/3 |  | 169.3 | 0.020 |
| VISli5 |  | 148.4 | 0.038 |
| VISpl4 |  | 22.2 | 0.075 |
| VISpl2/3 |  | 59.5 | 0.050 |
| VISpl5 |  | 54.4 | 0.083 |
| VISrl4 |  | 97.2 | 0.103 |
| VISrl2/3 |  | 105.4 | 0.082 |
| VISrl5 |  | 110.4 | 0.083 |

#### Conv kernel size for LGNv

The above methods allowed us to set kernel sizes for intracortical connections, but not subcortical ones. We set the kernel sizes for inputs to LGNv and VISp L4 according to receptive field sizes in these regions. Receptive fields are about 9 degrees in LGNv and 11 degrees in VISp (32). As mentioned in Section, *Cortical population constraints,* mouse visual acuity is approximately 1 pixel/degree, therefore we set kernel size of the connection from input to LGNv to 9×9. We then set the kernel size of the connection from LGNv to VISp to 3×3, such that the receptive field size for VISp was 11×11 pixels.

#### Summary tables

In Table 8, we summarize the calculated number of channels in each area (in parenthesis) and the kernel size for each Conv layer.

**Table 8.**
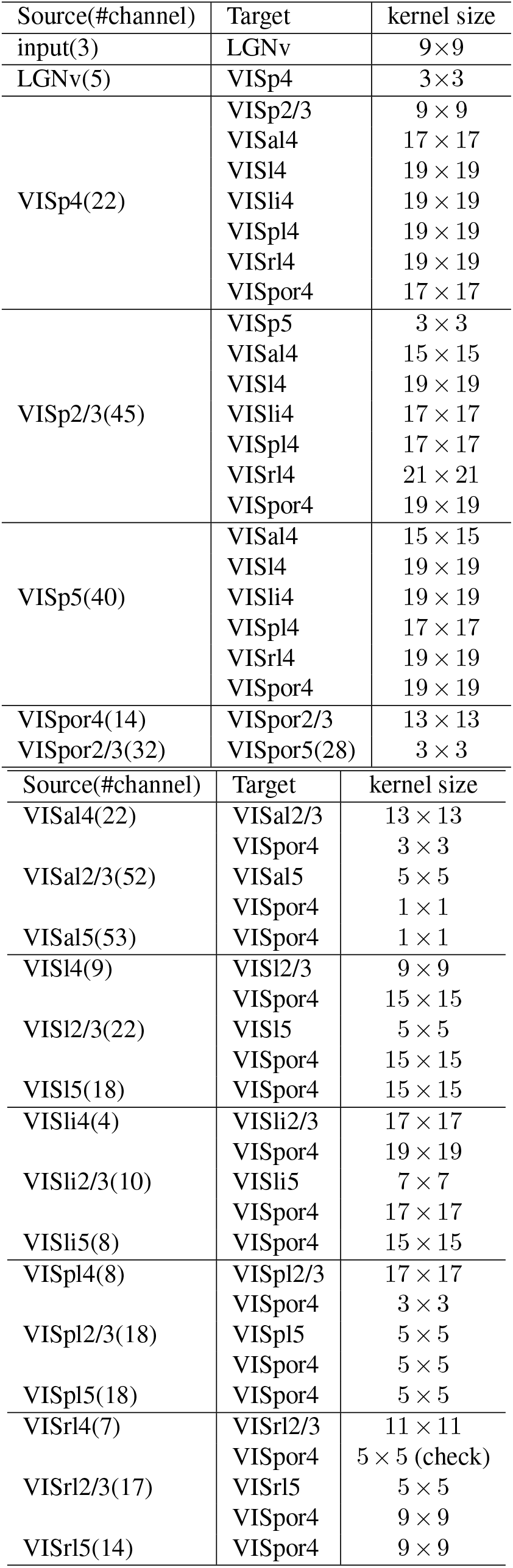
The calculated meta-parameters for the Conv layers.

The parameters used in the model based on biological sources and assumptions are summarized in Table 6 and the formulae for calculating the Conv layer meta-parameters are summarized in Table 9.

**Table 9.**
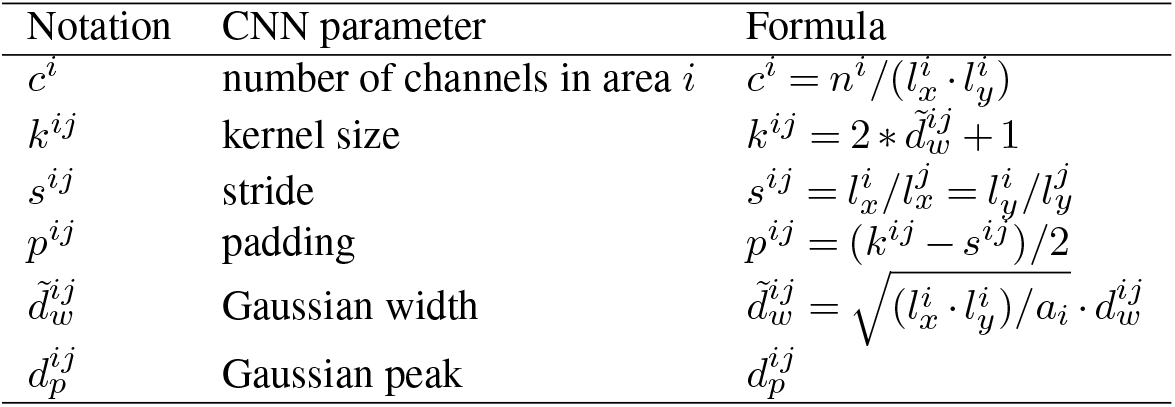
Meta-parameters for Conv layer connecting source area *i* to target area *j*

#### Implementation of MouseNet

We implemented the MouseNet structure in PyTorch. Each Conv layer was followed by a batch normalization layer and a ReLU nonlinearity linearity. For regions such as VISpor L4, which receive input from multiple Conv layers, the outputs of the Conv layers are summed before being fed into a batch normalization layer and a ReLU non-linnearity.

We have trained the MouseNet model on an image classification task, by adding a simple classifier. Specifically, the L5 output from all areas was reduced to 4×4 by an average pooling layer, then flattened, concatenated, and fed to a linear fully-connected layer which reduced the dimension to the number of classes of the task. The outputs were then transformed to probabilities by the softmax function and the cross-entropy loss was used to train on the image classification task. Analysis of the results is left for future work.

## Conclusions

Task-optimized deep networks show promise for brain modelling, because they are functionally sophisticated, and they often develop internal representations that overlap strongly with representations in the brain. Deep network architectures are loosely inspired by the brain, but there has been an extensive empirical exploration of the effects of architectural features in machine learning, independent of neuroscience. In parallel, a great deal has been learned about brain architectures, and mouse brain architecture has been particularly well characterized.

We have developed a deep network architecture that is consistent with a wide range of data on mouse visual cortex, including data from tract-tracing studies and studies of local connection statistics. While standard deep networks have provided useful points of comparison with neurobiological systems, more biologically realistic deep networks may enable more specific comparisons with the brain, such as comparisons between homologous groups of neurons, or modelling of specific lesions. By making our code publicly available, we hope to facilitate such work.

## ACKNOWLEDGEMENTS

We thank Tianqi Chen, Blake Richards, Shahab Bakhtiari, Graham Taylor for helpful discussions and suggestions on the manuscript. We thank Saskia de Vries, Hannah Choi, Kameron Decker Harris, Timothy Lillicrap, Julie Harris, and Severine Durand for helpful discussions. We thank the Allen Institute for Brain Science founder, Paul G. Allen, for his vision, encouragement, and support. We acknowledge the NIH Graduate training grant in neural computation and engineering (R90DA033461).

